# MR-pheWAS with stratification and interaction: Searching for the causal effects of smoking heaviness identified an effect on facial aging

**DOI:** 10.1101/441907

**Authors:** Louise A C Millard, Marcus R Munafò, Kate Tilling, Robyn E Wootton, George Davey Smith

## Abstract

Mendelian randomization (MR) is an established approach for estimating the causal effect of an environmental exposure on a downstream outcome. The gene x environment (GxE) study design can be used within an MR framework to determine whether MR estimates may be biased if the genetic instrument affects the outcome through pathways other than via the exposure of interest (known as horizontal pleiotropy). MR phenome-wide association studies (MR-pheWAS) search for the effects of an exposure, and a recently published tool (PHESANT) means that it is now possible to do this comprehensively, across thousands of traits in UK Biobank. In this study, we introduce the GxE MR-pheWAS approach, and search for the causal effects of smoking heaviness – stratifying on smoking status (ever versus never) – as an exemplar. If a genetic variant is associated with smoking heaviness (but not smoking initiation), and this variant affects an outcome (at least partially) via tobacco intake, we would expect the effect of the variant on the outcome to differ in ever versus never smokers. If this effect is entirely mediated by tobacco intake, we would expect to see an effect in ever smokers but not never smokers. We used PHESANT to search for the causal effects of smoking heaviness, instrumented by genetic variant rs16969968, among never and ever smokers respectively, in UK Biobank. We ranked results by: 1) strength of effect of rs16969968 among ever smokers, and 2) strength of interaction between ever and never smokers. We replicated previously established causal effects of smoking heaviness, including a detrimental effect on lung function and pulse rate. Novel results included a detrimental effect of heavier smoking on facial aging. We have demonstrated how GxE MR-pheWAS can be used to identify causal effects of an exposure, while simultaneously assessing the extent that results may be biased by horizontal pleiotropy.

**Author summary:** Mendelian randomization uses genetic variants associated with an exposure to investigate causality. For instance, a genetic variant that relates to how heavily a person smokes has been used to test whether smoking causally affects health outcomes. Mendelian randomization is biased if the genetic variant also affects the outcome via other pathways. We exploit additional information – that the effect of heavy smoking only occurs in people who actually smoke – to overcome this problem. By testing associations in ever and never smokers separately we can assess whether the genetic variant affects an outcome via smoking or another pathway. If the effect is entirely via smoking heaviness, we would expect to see an effect in ever but not never smokers, and this would suggest that smoking causally influences the outcome. Previous Mendelian randomization studies of smoking heaviness focused on specific outcomes – here we searched for the causal effects of smoking heaviness across over 18,000 traits. We identified previously established effects (e.g. a detrimental effect on lung function) and novel results including a detrimental effect of heavier smoking on facial aging. Our approach can be used to search for the causal effects of other exposures, where the exposure only occurs in known subsets of the population.

## Introduction

Mendelian randomization (MR) is an established approach that uses genetic variants as proxies for a modifiable exposure, to estimate the causal effect of the modifiable exposure on a downstream outcome (1). MR is often implemented within an instrumental variables (IV) framework, with a key assumption being that the genetic instrument affects the outcome solely through the exposure of interest. Horizontal pleiotropy (2), where the genetic instrument affects the outcome through pathways not via the exposure, invalidates this assumption. Whilst the absence of horizontal pleiotropy cannot be statistically tested, MR studies can include an investigation of evidence for a violation of this MR assumption (3). For instance, where several independent genetic instruments exist for an exposure, a difference in the effect estimates would indicate they may be acting through different pathways (4–6). The majority of these approaches are only applicable where there are several genetic variants that proxy for the exposure of interest (3,7).

A design in which the genetic variant is interacted with another variable can provide evidence regarding violation of the IV assumptions. The additional variable can be of many forms; not necessarily environmental. For example, an early study interacted a genetic variant related to alcohol with sex in a population where women drank very little, such that an effect of the variant on the outcome in women would suggest that at least some of the effect of the variant on the outcome is not acting through alcohol consumption (8). Here we report interaction with an environmental factor, and refer to the design henceforth as involving gene x environment (GxE) interaction (8–12). In contrast to traditional GxE studies that aim to identify genetic variants whose effects on an outcome are modified by an environmental factor (or vice-versa) (13), the aim here is to determine the extent that horizontal pleiotropy may be biasing estimated causal effects (12). To conduct a GxE MR study, a phenotype is chosen that stratifies the population into two (or more) groups, with different levels of the exposure – in the ideal case the exposure will manifest in one group but not in the other. If the effect of the genetic variant is (at least partly) via the exposure, the estimate of the direct test between the genetic variant and the outcome should vary in proportion to the degree the exposure manifests in each group. Importantly, the genetic instrument should not be associated with the phenotype used to stratify the sample, otherwise estimates may be biased due to conditioning on a collider (14).

The MR GxE approach has been used to investigate the effect of smoking heaviness on traits such as body mass index (BMI) and depression/anxiety (15–20). A causal effect of a genetic variant known to affect smoking heaviness should, if the effect is solely via tobacco intake, be seen in participants who are previous or current smokers, but not in participants who have never smoked. Thus, by stratifying on smoking status (ever versus never), we can assess if the smoking heaviness genetic variant is likely to be affecting an outcome via smoking heaviness, or some other pathway (or both). Figure 1 illustrates the three broad types of results of a GxE study on smoking heaviness: 1) no interaction between ever and never smokers, 2) a quantitative interaction, where associations are in a consistent direction but one is stronger than the other, or 3) a qualitative interaction, where the effect only occurs in one group or the estimates are in opposite directions, also known as a cross over effect (21). Both qualitative and quantitative interactions can provide evidence of a causal effect of the genetic variant via smoking heaviness. However, when an effect exists in never smokers the estimate in ever smokers may be comprised of both an effect of the genetic variant through smoking heaviness and an effect through another pathway.

**Figure 1:**
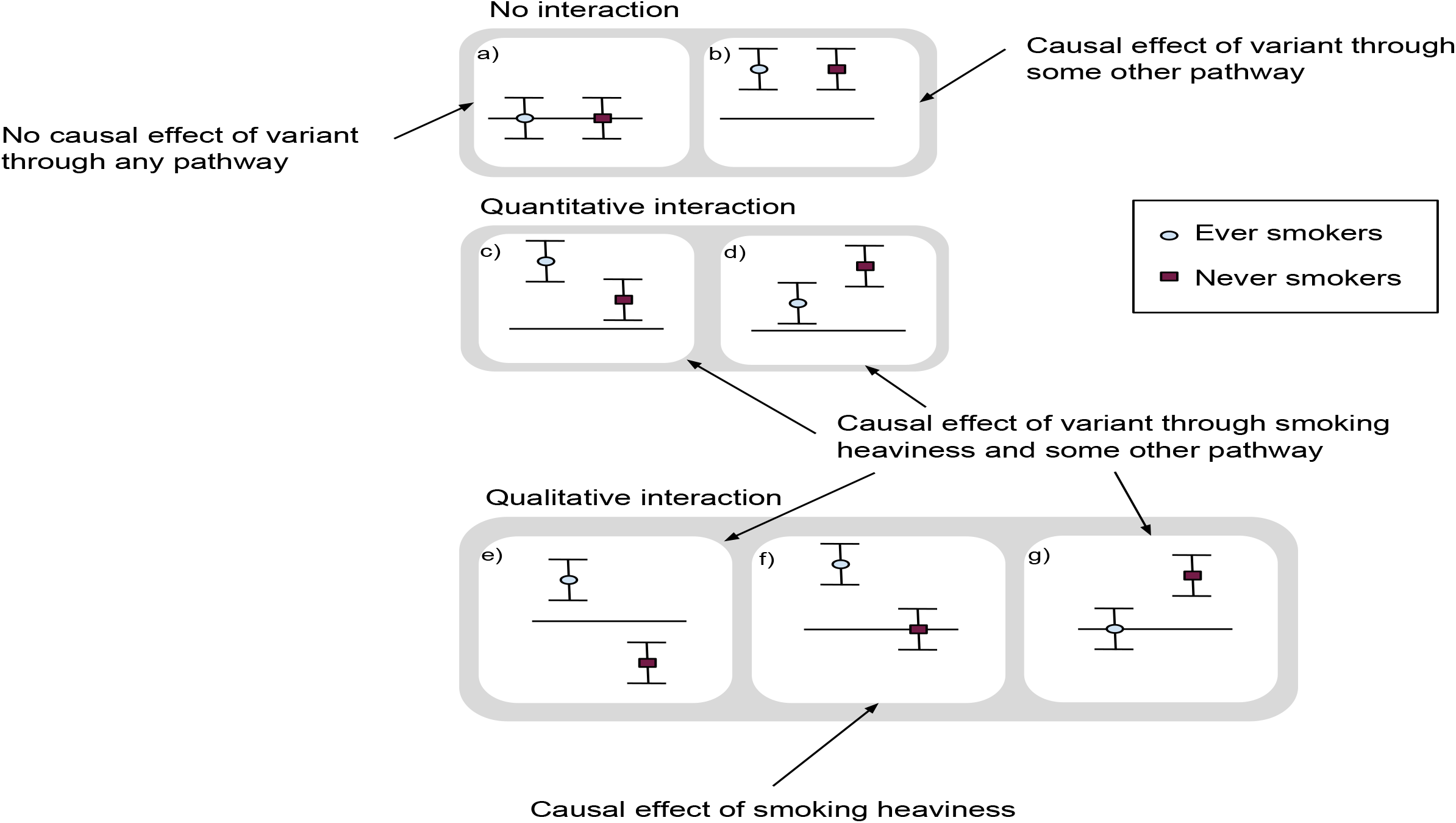
Illustration of possible GxE MR results and their interpretation

**Figure 2:**
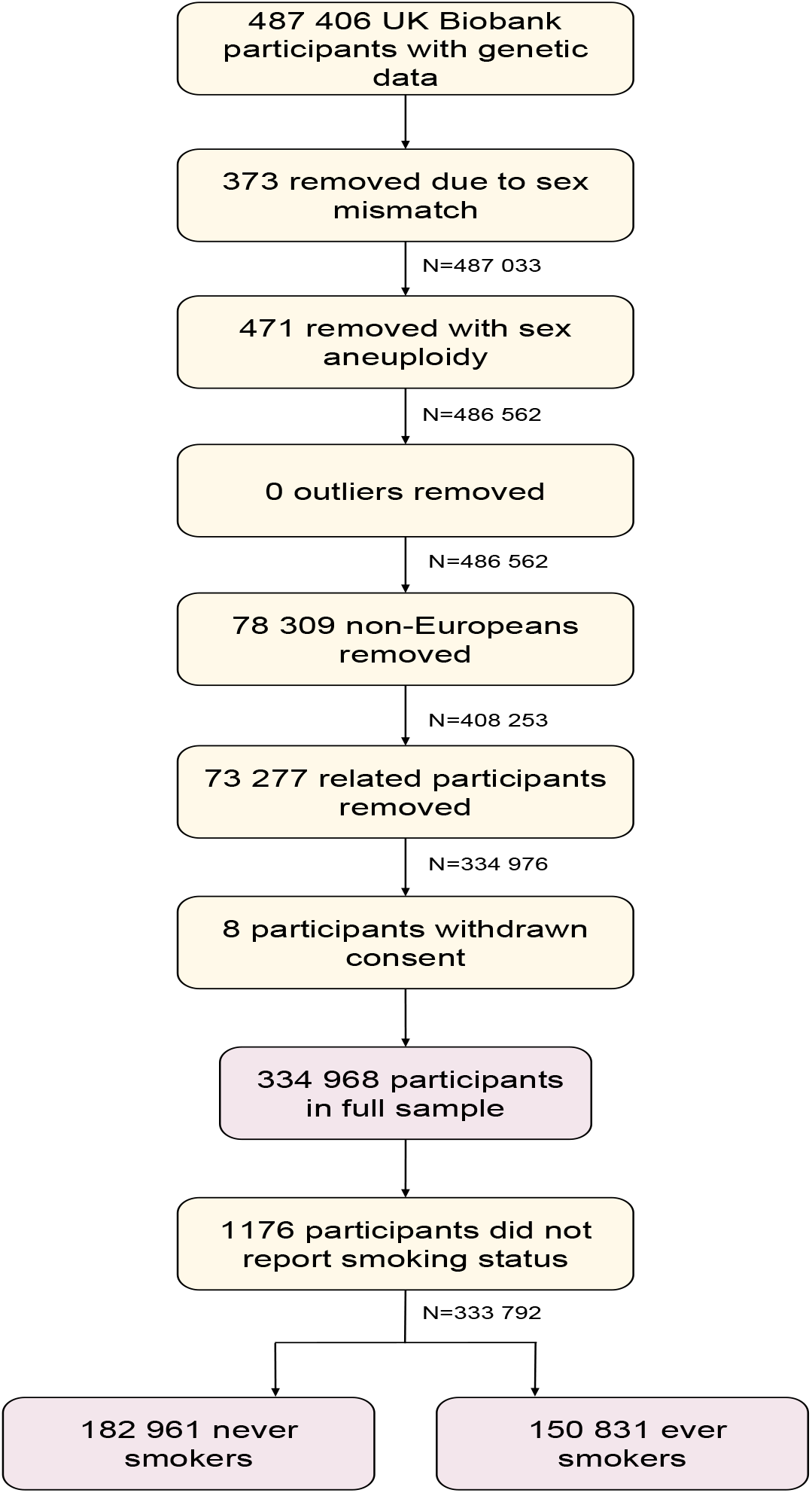
Participant flow diagram

While MR studies have, to date, been largely performed in a hypothesis-driven manner, it is now possible to perform hypothesis-free MR analyses using the MR phenome-wide association study (MR-pheWAS) approach (5,22). MR-pheWAS search for the causal effects of an exposure of interest, by testing the direct effect of a genetic instrument on a potentially large set of outcomes. Recently a tool for performing comprehensive phenome scans in UK Biobank– the Phenome Scan ANalysis Tool (PHESANT) – has been published (22). PHESANT allows researchers to search for the association of a trait of interest across over 18 000 phenotypes in UK Biobank, and is the first freely-available tool for performing comprehensive phenome scans. Here we introduce the GxE MR-pheWAS approach – searching for the causal effects of an exposure, while simultaneously providing evidence of the degree to which an estimated effect may be biased due to horizontal pleiotropy. GxE MR-pheWAS can be performed using PHESANT. We perform a proof of principle study, searching for the effects of smoking heaviness.

## Results

Each additional smoking-increasing allele of rs16969968 was associated with a 1.21 [95% confidence interval (CI): 1.19, 1.23] higher odds of being in a higher smoking heaviness category, after adjusting for age, sex and the first 10 genetic principal components, and a 0.98 [95% CI: 0.97, 0.99] lower odds of being an ever (vs never) smoker.

### Results of GxE MR-pheWAS analysis

#### Identified main effects from MR-pheWAS in ever smokers

The results of our MR-pheWAS among ever smokers includes 18 514 tests ranked by P value of the estimated effect on each outcome, given in S1 File (S1 Fig in S1 Text shows the number of fields reaching each stage of the PHESANT pipeline). A QQ plot is given in Figure 3a. We identified 70 results at a false discovery rate of 5% (using a P value threshold of 0.05×70/18 514=1.89×10^-4^), given in Figure 4 and see Table B in S1 Text, and of these, 30 results had a P value lower than a stringent Bonferroni corrected threshold of 2.70×10^-6^ (0.05/18 514). In contrast, we identified 8 results at a false discovery rate of 5% among never smokers (see Table C in S1 Text), 47 results among our full sample (see Table D in S1 Text), and 9 results using a two-step approach (identifying the outcomes associated using the full sample then ranking by interaction strength among this subset; see Table F in S1 Text).

**Figure 3:**
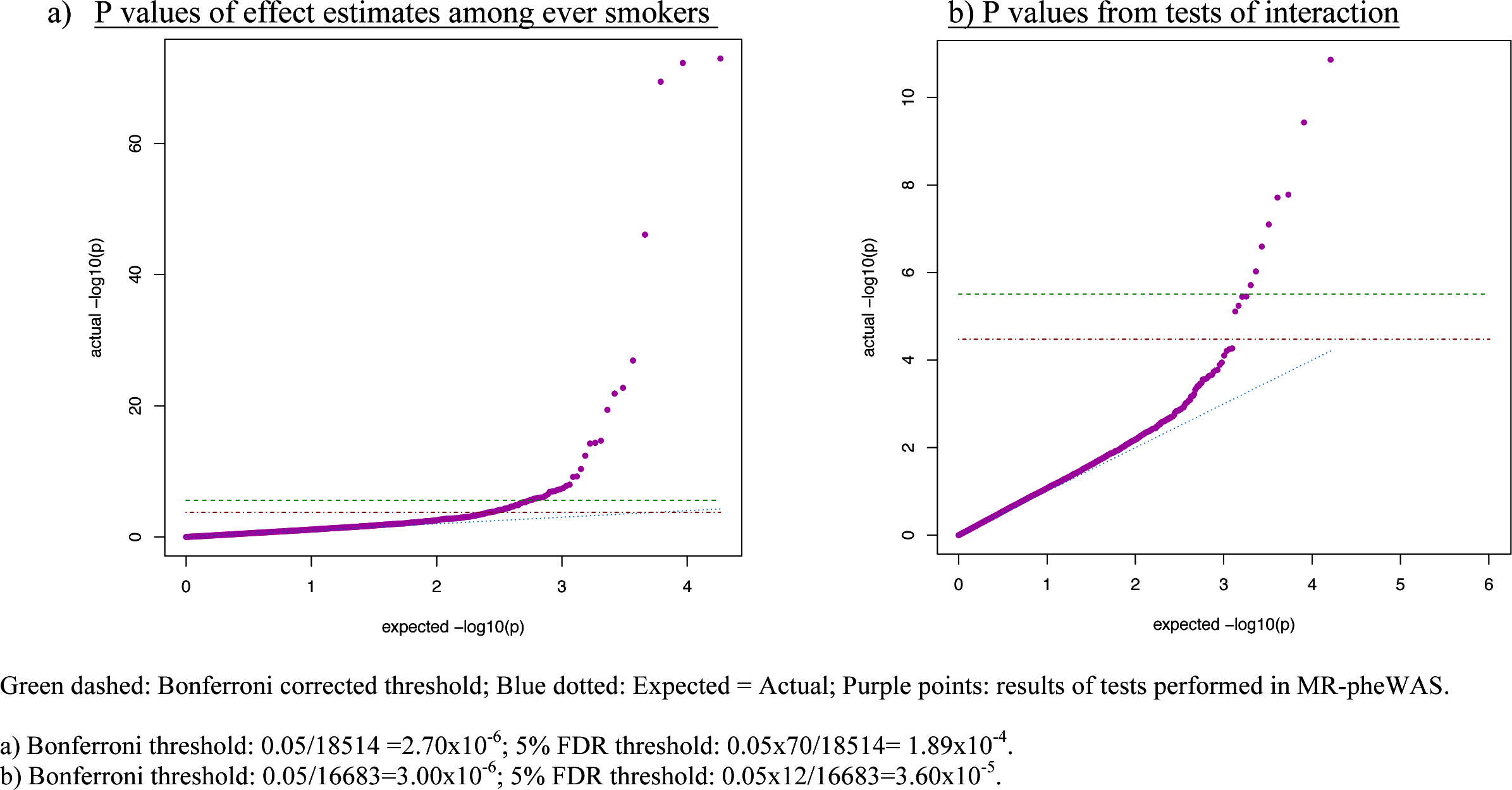
QQ plots of PHESANT MR-pheWAS results

**Figure 4:**
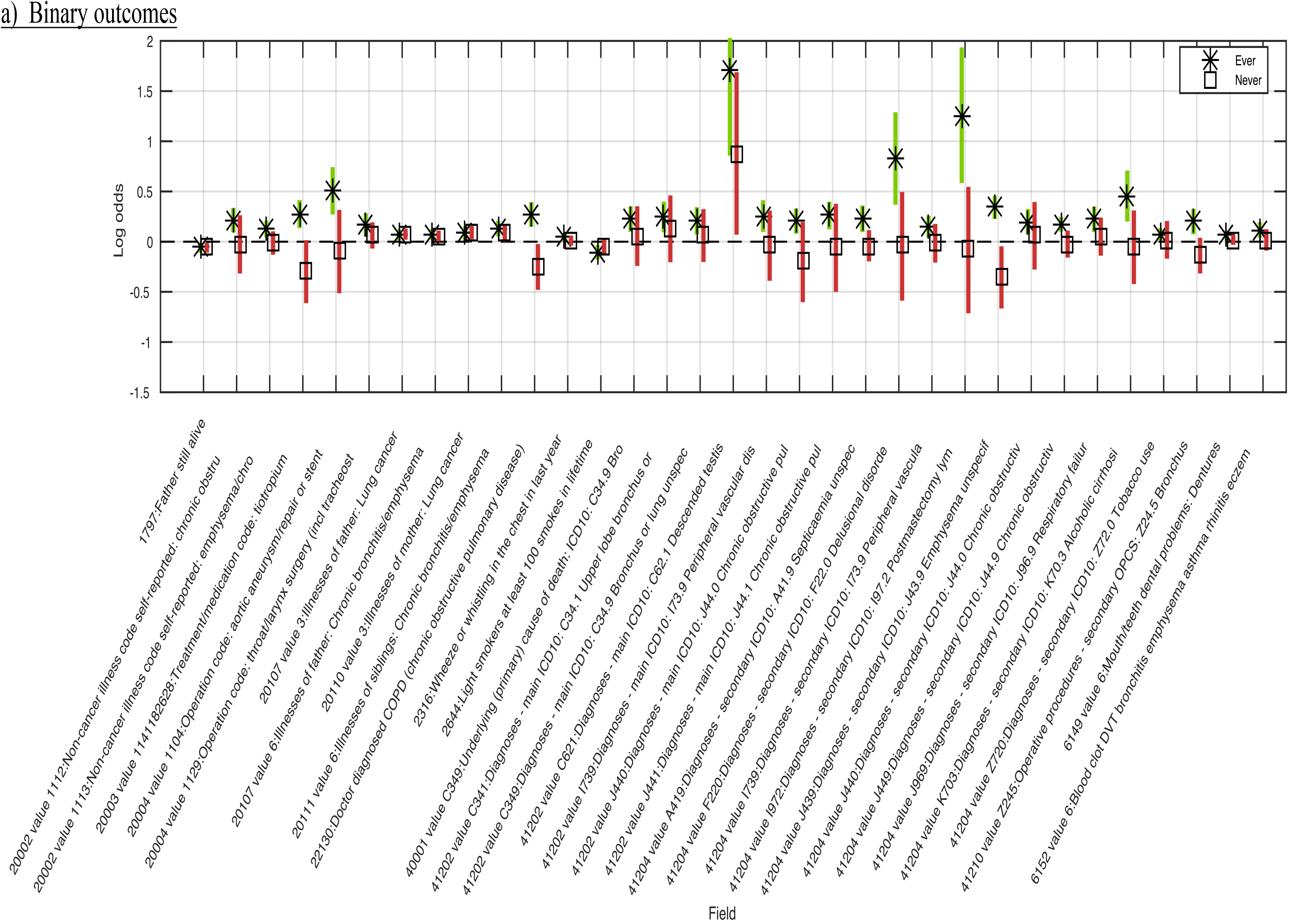
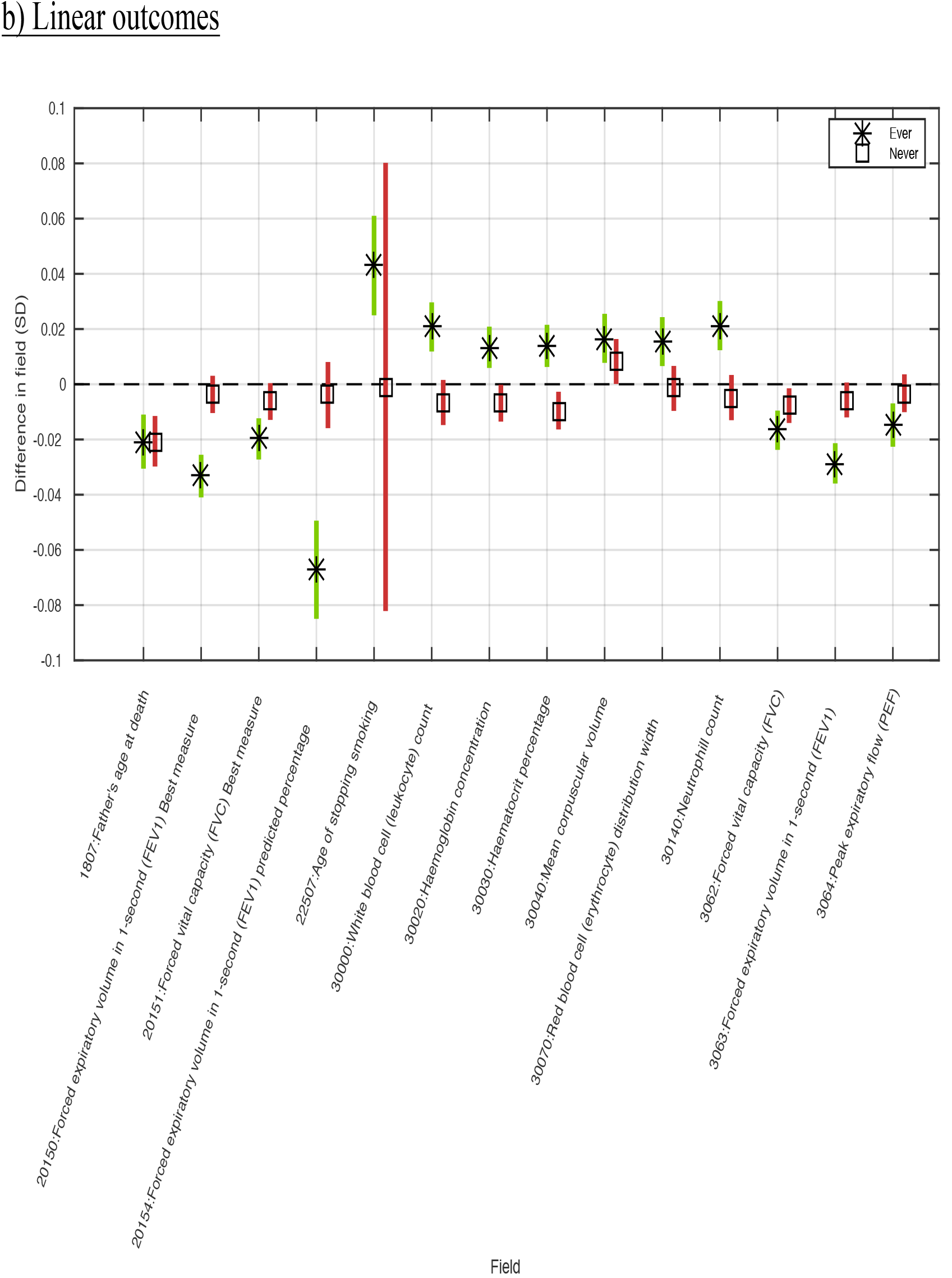
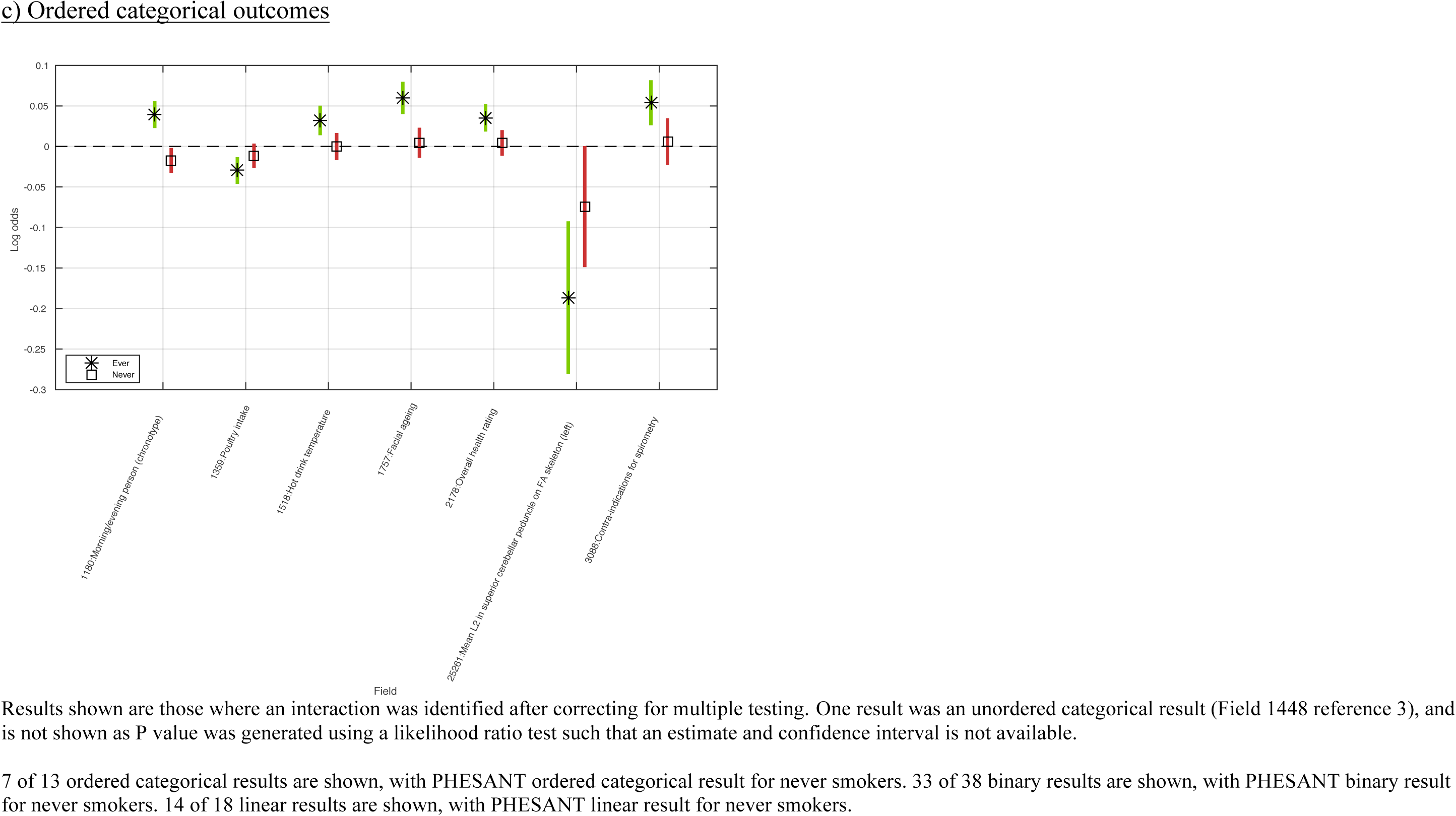
Identified main effects from MR-pheWAS in ever smokers

#### Identified interactions from MR-pheWAS in ever smokers versus never smokers

Of the 16 683 interactions tested, we identified 8 results with a P value lower than a stringent Bonferroni corrected threshold of 3.00×10^-6^ (0.05/16 683), where an increase in rs16969968 smoking-heaviness increasing allele dosage was associated with worse lung function (3 phenotypes), and less likelihood of being a morning person, in ever compared with never smokers. We found a further 4 results at a false discovery rate of 5% (using a P value threshold of 0.05×12/16 683=3.60×10^-5^) (see Table E in S1 Text), where an increase in rs16969968 smoking-heaviness increasing allele dosage was associated with a higher risk of chronic obstructive pulmonary disease, emphysema and cancer diagnoses and more facial aging, in ever compared with never smokers. A QQ plot is given in Figure 3b.

The results identified when ranking by our P value for interaction were a subset of those identified when ranking by the P value of the main effects among ever smokers, except for a binary outcome describing whether the participant has had a keratinizing squamous cell carcinoma (field ID [FID]=40011, value 8071). A higher rs16969968 smoking-heaviness increasing allele dosage was associated with a higher risk of keratinizing squamous cell carcinoma among ever smokers, but a lower risk among never smokers. Overlap of results across all our MR-pheWAS is shown in S2 Fig in S1 Text.

#### Detailed follow-up of potentially novel results

Our identified results included a detrimental effect of heavier smoking on facial aging (FID=1757), where a higher genetic predisposition to heavier smoking was associated with an increased risk of looking older (as perceived by others) relative to your age. Estimates of association are shown in Table 1. Among ever smokers each additional smoking-increasing allele of rs16969968 was associated with a 1.062 [95% CI: 1.043, 1.081] higher odds of reporting an older facial aging category, and a 1.004 [95% CI: 0.988, 1.021] higher odds among never smokers (interaction P=7.72×10^-6^). Our sensitivity analyses adjusting for age, sex and the first 40 principal components were consistent with the results of our main analyses.

**Table 1:** Results of associations of genetic instruments with facial aging outcome

| Analysis | Sample | N | Odds ratio | Interaction P value<sup>3</sup> |
| --- | --- | --- | --- | --- |
| Associations of rs16969968 on facial aging<sup>1</sup> |  |  |  |  |
| Main analysis | Ever smokers | 137,869 | 1.062 [1.043, 1.081] | 7.72x10<sup>-6</sup> |
|  | Never smokers | 167,781 | 1.004 [0.988, 1.021] |  |
| Sensitivity analysis | Ever smokers | 137,869 | 1.062 [1.043, 1.081] | 7.46x10<sup>-6</sup> |
|  | Never smokers | 167,781 | 1.004 [0.988, 1.021] |  |
| Estimates of causal effect of lifetime smoking on facial aging<sup>2</sup> |  |  |  |  |
| Main analysis | Ever and never smokers | 305,662 | 1.284 [1.085, 1.521] | NA |
| Sensitivity analysis |  |  | 1.286 [1.080, 1.531] | NA |

Main analysis: Adjusted for age, sex and first 10 genetic principal components. Sensitivity Analysis: Adjusted for age, sex and first 40 genetic principal components.
<sup>1</sup> Direct test of association between smoking heaviness SNP and facial aging, in ever and never smokers separately. Estimates are the change of odds of reporting looking 'older than you are' versus looking 'younger than you are' or 'about the same', or looking 'older than you are' or 'about the same' versus 'younger than you are', for each additional smoking-increasing allele of rs16969968.
<sup>2</sup> Two stage IV probit regression. Estimates are the change of odds of reporting facial aging category 'older than you are' for a 1 SD increase in lifetime smoking score. Calculated by taking the exponent of 1.6 times the probit estimate (43).
<sup>3</sup> Interaction P value generated using meta regression (`metan` command in Stata).

We attempted to replicate this effect using another instrument for smoking behavior, which captures lifetime smoking exposure as opposed to just smoking heaviness. A 1 standard deviation (SD) higher lifetime smoking genetic instrument was associated with a 0.104 [95% confidence interval (CI): 0.101, 0.107] SD higher lifetime smoking score in our sample, after adjusting for age, sex and the first 10 genetic principal components. We estimated that a 1SD higher lifetime smoking score caused a 1.284 [95% CI: 1.085, 1.521] higher odds of reporting that people say you look older than you are, after adjusting for age, sex and the first 10 genetic principal components.

#### Testing the impact of collider bias

Our conclusions about the causal effect of smoking heaviness on our outcomes may be affected by collider bias (14). This is because we stratified on smoking status, and we found an effect of the smoking heaviness genetic variant on smoking status. If any cause of an outcome also causes smoking status, then an association may be induced between the smoking heaviness genetic variant and that outcome (S3 Fig in S1 Text). If there is in truth an association between the genetic variant for smoking heaviness and the outcome, then it could be estimated with bias in this scenario. We performed a simulation to determine the degree of confounding needed to induce an association between the genetic variant and our identified facial aging phenotype, given there is no true causal effect of the SNP on the outcome. Further details of our simulation approach are given in Section S1 in S1 Text. Our simulation indicated that any collider bias due to the effect of the genetic variant on smoking status is likely to have had a negligible impact on the estimated effect of the genetic variant on facial aging (S4 Fig in S1 Text), given the same sample size and numbers of ever and never smokers as our UK Biobank sample. For example, assuming no effect of the SNP on the outcome, and given both a confounder that is the only cause of facial aging, and large effect of the confounder on smoking status (OR=10), the estimated effect of the SNP on the facial aging in ever and never smokers are consistent and their confidence intervals both include the null, i.e. the degree of bias is not sufficient to (incorrectly) suggest an effect of the SNP on facial aging via either smoking heaviness or some other path.

## Discussion

In this study, we searched for causal effects of smoking heaviness, using the PHESANT package to perform a GxE MR-pheWAS, by estimating the association of genetic variant rs16969968 with each outcome while restricting to ever and never smokers, respectively. We used two approaches to identify the causal effects of smoking heaviness from our PHESANT results. The first, ranking results based on the strength of the effect of rs16969968 among ever smokers, identifies causal effects under the assumption that the combined effect through smoking heaviness and any pleiotropic effect is not null. For instance, given a positive effect of a genetic variant on an outcome via smoking heaviness, and an equal but opposite effect via horizontal pleiotropy, then the effect of the variant would appear null in ever smokers and negative in never smokers, and hence this outcome would not be identified when ranking by the effect among ever smokers. The second approach, ranking results by the strength of interaction between ever and never smokers, is commensurate with our aims, as a genetic variant that affects an outcome (at least in part) through smoking heaviness will exhibit a different effect size on this outcome, among ever and never smokers, respectively. However, this approach lacks statistical power, especially when combined with the need to correct for the multiple tests performed in an MR-pheWAS.

Our GxE MR-pheWAS of smoking heaviness identified 70 results when ranking on strength of effect in ever smokers, and 8 when ranking on interaction strength. Of the 8 identified in the latter, 7 of these were also identified in the former. Furthermore, the majority of results identified when ranking on the effect in ever smokers were qualitative, with an effect seen among ever smokers but not among never smokers (type F in Figure 1). Our results confirmed several established or previously reported causal effects of smoking heaviness. For example, our findings suggested higher smoking heaviness was associated with worse lung function (FIDs={3063, 20150, 20151}) (23), higher resting heart rate (FID=102) (24,25), higher risk of chronic obstructive pulmonary disease (FID=22130) (23) and lung cancer (FID=40001 value C349) (26), and lower odds of being a morning person (FID=1180) (27).

Our PHESANT-estimated results identified a weak negative effect of rs16969968 on BMI in ever smokers (that did not meet the 5% FDR threshold). Previous MR studies have also identified an effect of smoking heaviness on BMI, where additional smoking-increasing alleles was associated with a lower BMI among current smokers (10,15,19). Since we used ever smokers rather than current smokers our weaker association may be because the effect on BMI weakens over time after smoking cessation (10). Furthermore, while previous studies found an effect of rs16969968 on BMI in never smokers (10,15,19), we found little evidence of an association in UK Biobank.

We identified other novel results, including a detrimental effect on facial aging. We estimated that a 1SD increase in lifetime smoking causes a 1.28 [95% CI: 1.08, 1.52] higher odds of reporting that others say you look older than you are. A 1SD increase in lifetime smoking is, for example, equivalent to being a current smoker who has smoked 5 cigarettes per day for 12 years, or a former smoker who smoked 5 cigarettes per day for 21 years but stopped smoking 10 years ago, rather than a never smoker. This identified association should be further investigated and replicated in an independent sample; it may reflect a true causal effect of smoking heaviness, may be due to chance, or may arise because the genetic variants have horizontal pleiotropic effects and are thus invalid instruments for smoking heaviness or lifetime smoking. However, we examined the extent to which pleiotropy might be biasing our smoking heaviness results, by estimating the effect of the smoking heaviness genetic variant among never smokers, and found little evidence of an association. Furthermore, another recent study assessed perceptions of facial attractiveness using a two-alternative forced choice design where participants were randomly shown prototypical faces for smokers and non-smokers and asked to select the most attractive, and found that smoking faces were deemed less attractive (28). Our study using MR is likely to have different sources of potential biases such that triangulation of these results adds further evidence of the detrimental effect of smoking on facial appearance in general (29,30).

Our results include examples of the “case-control by proxy” study design, where participants with relatives who are cases for a given phenotype are used as ‘proxy’ cases and those with relatives who are controls are used as ‘proxy’ controls (31). For example, we identified associations with parental risk of smoking related diseases, such as lung cancer and emphysema, with consistent estimates in ever and never smoking subsamples, likely due to the shared genetic risk of the UK Biobank participant with their parents. Although we have not identified examples in our top results, associations may also reflect an effect of the parental genotype of UK Biobank participants on the phenotype of the UK Biobank participant. For example, maternal genotype could influence participant’s phenotype via intrauterine exposure to maternal smoking.

Our comparison of results across samples demonstrates the value of stratifying to detect potential causal effects. Of the 70 results identified in ever smokers, only 36 were identified in our MR-pheWAS using the full sample. While the results of our MR-pheWAS in the full sample included associations that were not identified in our MR-pheWAS in ever smokers, such as risk of operative procedures on the tarsometatarsal joint and eye problems, these associations may be due to pleiotropy (or chance) rather than a causal effect of the smoking heaviness variant through smoking status, as associations were consistent in ever and never smokers.

There are some limitations of this work that should be considered. First, due to the automated processing of PHESANT where pre-processing of an outcome may depend on the number of participants with each outcome value, a different statistical test is used for some outcomes (e.g. linear versus ordered logistic regression), for ever versus never smokers, such that tests of interaction between these could not be performed. Second, while ranking by interaction strength between ever and never smokers would be the preferred approach to identify causal effects of smoking heaviness (i.e. outcomes where the effect of the genetic variant differs among ever versus never smokers), this lacked power and identified only a small number of results. However, ranking using the strength of association among ever smokers has better power and, although in theory some results may be missed, in practice we identified many potentially interesting novel results with this method, including all but one of the results based on interaction strength.

Third, UK Biobank is a highly selected sample of the UK population, having a response rate of 5.5% (32). For example, UK Biobank participants are, on average, less likely to smoke, and have less self-reported health conditions, compared with the general population (33). Among smokers, the difference between smoking heaviness in UK Biobank compared with the general population varies by age and sex. For instance, smokers aged 45-54 years in UK Biobank, and women smokers aged 55-64 years in UK Biobank on average smoke more heavily than those in the general population, but this difference was not seen among male smokers age 55-64. If these differences are due to an effect of smoking heaviness on selection into the study, then our estimates may be biased (34). Also, if selection is additionally dependent on a given outcome, then associations may be biased by a particular form of collider bias – selection-induced collider bias (14). In general, collider bias may occur when two variables (A and B) affect a third variable and this third variable (C) is conditioned upon in analyses. Selection induced collider bias may occur when variable C represents whether a person is selected into the sample (i.e. variables A and B both affect participation in the study). Hence, estimates of association between two phenotypes – such as our smoking heaviness genetic variant and a given outcome in our study – can be biased, if inclusion in the study is affected by both phenotypes.

Fourth, we found an association between rs16969968 and smoking status (ever versus never). This means that associations may be biased by selection induced collider bias, because we stratify our sample on smoking status. While our simulations indicated that any collider bias due to the SNP effect on smoking status would have a negligible impact on the estimated effect of the SNP on a given outcome, it is possible that this has increased our type 1 error rate across the large number of tests performed in our MR-pheWAS.

Fifth, the estimates used to identify interactions are of the direct association of the genetic variant with the outcome, and so are not estimates of the causal effect of smoking heaviness. We cannot follow-up results using a formal IV analysis because there is no accurate measure of tobacco exposure, and using poor measures such as cigarettes smoked may give biased associations (7). For other exposures that are measured accurately, GxE interactions can be used to estimate the size of causal effects in the presence of pleiotropy (35).

Sixth, it is possible that reporting bias of smoking status may have biased associations. For instance, if some ‘ever’ smokers reported that they have never smoked, then for outcomes affected by rs16969968 via smoking heaviness the estimate in never smokers would be biased towards that in ever smokers. This would bias estimates of interaction between ever and never smokers towards the null. Furthermore, if the effect of smoking heaviness is transient then our interaction estimates may also be biased towards the null, because previous smokers are assigned to the ever smoker group but the effect may no longer be present. In this case, testing the interaction between current and never smokers may be more appropriate.

Seventh, due to the hypothesis-free nature of a phenome scan, results generated in this way require careful consideration and follow-up. For example, our MR-pheWAS identified a potentially interesting association of rs16969968 with risk of diagnosis of the International Classification of Diseases (ICD) code ‘Descended testis’ (field 41202 value C621). This association was found in both ever and never smokers, hence we initially considered whether this may reflect an effect of parental genotype on offspring phenotype. However, further inspection revealed that this result is misleading for two reasons. First, this ICD code (C62.1) is a subcategory of ‘Malignant neoplasm of testis’ [C62]), hence pertains specifically to cancerous descended testis. Second, PHESANT deals with ICD fields by generating a binary variable for each ICD code, assigning all participants with the code as TRUE and all other participants as FALSE (i.e. assuming no missingness across all participants). This is not appropriate for sex specific codes, where analyses should be restricted to a particular sex. We further investigated this result by restricting to male participants, and testing the effect of rs16969968 on: 1) ‘malignant neoplasm of testis, descended’ and 2) ‘malignant neoplasm of testis, unspecified’. The latter serves as a replication for the former, under the assumption that the proportion of participants with descended versus undescended testis in the ‘unspecified’ group is the same as the ICD codes (C62.1 versus C62.0) where this is known (i.e. the majority of the unspecified group are descended; the ratio in UK Biobank is 1:16). While the positive association with the ‘descended’ group remains in both ever and never smokers (N_descended=26), we find little evidence of an association in the unspecified group (N_unspecified=199), suggesting that the association with the particular data field is due to chance.

We used the freely available PHESANT package to search for causal effects across thousands of outcomes. We have shown that the GxE MR-pheWAS is an effective approach to search for the causal effects of an exposure using observational data, when the degree to which an exposure manifests is different across known subsets of the population. Other potential applications include searching for the causal effects of alcohol intake (stratified by never versus ever drinkers), cannabis use (stratified by never versus ever used) and milk consumption (stratified by ever or never drinking milk). The approach could also be applied to qualitative gene by gene interactions. Our study of smoking heaviness serves as a model for future studies seeking to search for the causal effects of an exposure using GxE MR-pheWAS.

## Materials and methods

### Study population

UK Biobank is a prospective cohort of 503 325 men and women in the UK aged between 37–73 years (99.5% were between 40 and 69 years) (36). This cohort includes a large and diverse range of data from blood, urine and saliva samples and health and lifestyle questionnaires (37).

Of the 487 406 participants with genetic data, we removed 373 with genetic sex different to reported sex, and 471 with sex chromosome aneuploidy (identified as putatively carrying sex chromosome configurations that are not either XX or XY). We found no outliers in heterozygosity and missing rates, which would indicate poor quality of the genotypes. We removed 78 309 participants not of white British ancestry (38). We removed 73 277 participants who were identified as being related, having a kinship coefficient denoting a third degree (or closer) relatedness (38). We removed 8 individuals with withdrawn consent, giving a sample of 334 968 participants (we refer to as our full sample). Of these, 182 961 and 150 831 reported being never and ever (comprising previous and current) smokers, respectively. A participant flow diagram is given in Figure 2.

### Genetic variant for smoking heaviness

We use genetic variant rs16969968 within the *CHRNA5* gene, as an instrument for smoking heaviness (16), coded as the number of smoking heaviness increasing alleles.

### Outcomes

The UK Biobank data showcase allows researchers to identify variables based on the field type (http://biobank.ctsu.ox.ac.uk/showcase/list.cgi). At the time of data download there were 2761 fields of the following types: integer, continuous, categorical (single) and categorical (multiple).

We excluded 74 fields *a priori* (see Table A in S1 Text) for the following reasons: 1 field denoting the assessment centre; 2 fields described by UK Biobank as ‘polymorphic’, containing values with mixed data types; 7 fields that, although listed in the data showcase, were not currently available; 17 genetic descriptor fields, 1 sex field, 4 age fields, 17 fields describing the assessment centre environment, 4 data processing indicators, and 21 categorical (single) fields with more than one value recorded per person.

This resulted in a set of 2687 UK Biobank fields (347 integer, 1392 continuous, 836 categorical [single] and 112 categorical [multiple]), referred to hereafter as the outcome dataset (because they are tested as an outcome irrespective of whether this is biologically plausible).

### Smoking phenotypes

Smoking status was self-reported via a questionnaire at the UK Biobank assessment centre. Participants were asked to report whether they smoked previously, currently, or whether they had never smoked. We created a binary variable denoting ever versus never smokers by grouping former and current smokers.

Smoking heaviness was derived from the number of cigarettes smoked per day, which was asked via the same questionnaire, to those who reported being a previous or current smoker. We categorised the number of cigarettes into four bands: 0-10, 11-20, 21-30 and 31+.

### Covariates

We include age and sex as covariates in our models to reduce the variation in our outcomes. Age when participants attended the UK Biobank assessment centre was derived from their date of birth and the date of their assessment centre visit. Sex was self-reported during the touchscreen questionnaire (and validated using the genome-wide data). We adjusted for the first 10 genetic principal components to control for confounding via population stratification. Genetic variants are set at conception, and after conception they cannot be affected by traditional confounding factors, therefore we did not adjust for any further covariates.

### Statistical methods

#### Smoking heaviness genetic variant and self-reported smoking heaviness association

We tested the association of rs16969968 with smoking heaviness using ordered logistic regression (`ologit` Stata command), adjusting for covariates as described above.

#### PHESANT MR-pheWAS

We searched for the causal effects of smoking heaviness, within three subsamples of UK Biobank participants: 1) ever smokers, 2) never smokers, and 3) our full sample. For each subsample, we tested the direct association of rs16969968 with each of the outcome variables using the PHESANT package (version 0.15). A description of PHESANT’s automated rule-based method is given in detail elsewhere (22). In brief, the decision rules start with the variable field type and use rules to categorize each variable as one of four data types: continuous, ordered categorical, unordered categorical or binary. Variables with the continuous and integer field type are usually assigned to the continuous data type, but some are assigned to ordered categorical if, for instance, there are only a few distinct values. Variables of the categorical (single) field type are assigned to either the binary, ordered categorical or unordered categorical, depending on whether the field has two distinct values, or has been specified as ordered or unordered in the PHESANT setup files. Variables of the categorical (multiple) field type are converted to a set of binary variables, one for each value in the categorical (multiple) fields.

PHESANT estimates the univariate association of rs16969968 with each outcome variable. The rs16969968 SNP and outcome are the independent (exposure) and dependent (outcome) variables in the regression model, respectively. Outcome variables with continuous, binary, ordered categorical and unordered categorical data types, were tested using linear, logistic, ordered logistic, and multinomial logistic regression, respectively. Prior to testing, an inverse normal rank transform was applied to variables of the continuous data type, to ensure they were normally distributed. All analyses were adjusted for covariates as described above.

#### Identifying results based on strength of effect in ever smokers

Under the assumption that an effect of the rs16969968 variant operates through smoking heaviness, associations that reflect causal effects of smoking heaviness should be seen only in ever smokers. Hence, to identify causal effects we ranked outcomes by P value of the estimated effects of rs16969968 within the ever subsample only. We corrected for multiple testing by controlling for the expected proportion of false positive results. We identified the largest rank position with a P value less than *P_threshold_*= 0.05×rank/n, where *n* is the number of total number of tests in the phenome scan. *P_threshold_* is the P value threshold resulting in a false discovery rate of 5% (39). We also calculated a highly stringent Bonferroni corrected P value calculated by multiplying each P value by the number of tests performed, which assumes each test is independent. We examined the degree to which pleiotropy may be biasing results by viewing these estimates alongside the estimates among never smokers. We also identified top results in our never smoker and full samples, to compare the sets of results identified using each sample.

#### Identifying results based on strength of interaction of ever versus never smokers

PHESANTs rule-based method decides how to appropriately test associations and in some cases, for a given outcome, different decisions are taken for the ever versus never subsets of UK Biobank. For this reason, we first identified the subset of outcomes, where PHESANT used the same type of regression to test the association, among ever and never smokers. For this subset, we determined the strength of interaction between ever versus never smokers using meta regression (`metan` command in Stata), and ranked the results by the P value of the generated Q-test statistic – a measure of the heterogeneity across subgroups. We tested interactions of binary and ordered categorical results in terms of the log odds (i.e. the interactions are assumed to be multiplicative). To identify potential causal effects, we used both a Bonferroni and 5% FDR threshold, as described above.

It may be reasonable to assume that rs16969968 affects most outcomes only via smoking heaviness (i.e. there is no horizontal pleiotropy), such that an association between rs16969968 and the outcome in the whole sample would indicate an effect of smoking heaviness on this outcome. We perform a two-step approach to identify interactions, similar to an approach for identifying gene-environment interactions proposed previously (40). First, we rank outcomes by the strength of association in the whole sample and used a 5% FDR threshold to identify results to take forward to step two. Second, we rank these results by the strength of interaction between ever and never smokers, ranking by the P value of the Q-test statistic. This approach is valid because the strength of association in the whole sample is independent of the strength of the interaction between ever and never smokers.

#### Follow-up analyses of identified associations

We identified an association with a facial aging phenotype. Participants were asked ‘do people say you look:’ and asked to select either ‘younger than you are’, ‘about your age’ or ‘older than you are’. It is possible that the PHESANT automated approach made inappropriate decisions in its analysis, hence we re-examined this association to ensure it is not erroneous. We estimated the effect of rs16969968 on facial aging, using ordered logistic regression (ologit Stata command), among ever and never smokers respectively, adjusting for covariates as described above. We also adjusted for age, sex and the first 40 genetic principal components, as a sensitivity analysis.

We derived a measure of lifetime smoking that incorporates smoking heaviness, duration and time since cessation into a single measure (41) called the lifetime smoking index as described previously (42). We generated an lifetime smoking genetic instrument using the 124 identified SNPs from a recent GWAS of the lifetime smoking index (42). As this GWAS was also conducted in UK Biobank, we calculated our lifetime smoking genetic instrument as the sum of the lifetime smoking index increasing alleles (i.e. we did not weight by the effect size). We derived a binary measure of facial aging representing whether ‘people say you look older than you are’ or not, by combining the ‘younger than your age’ and ‘about your age’ categories. We determined the strength of our lifetime smoking genetic score as an instrument for lifetime smoking index using linear regression, adjusting for age, sex and the first 10 genetic principal components. We generated confidence intervals using bootstrapping (with 1000 bootstraps) as the residuals of this regression (due to non-normality of the lifetime smoking index) were not normally distributed. We estimated the causal effect of lifetime smoking on facial aging, using IV probit regression. Again, we generated confidence intervals using bootstrapping (with 1000 bootstraps) as the residuals of the first stage were not normally distributed. We take the exponent of 1.6 times the estimates, to approximate the association in terms of the change of odds (43).

Analyses are performed in R version 3.2.4 ATLAS, Matlab r2015a or Stata version 14, and code is available at [https://github.com/MRCIEU/PHESANT-MR-pheWAS-smoking]. Git tag v0.1 corresponds to the version presented here.

## Acknowledgements

This research has been conducted using the UK Biobank Resource under Application Number 16729.

## Author contributions

LACM contributed to the design of the study, performed the analyses, drafted the initial manuscript, reviewed and revised the manuscript and approved the final version of the manuscript as submitted. MRM contributed to the design of the study, reviewed and revised the manuscript and approved the final version of the manuscript as submitted. KT contributed to the design of the study, reviewed and revised the manuscript and approved the final version of the manuscript as submitted. GDS conceptualized and designed the study, reviewed and revised the manuscript and approved the final version of the manuscript as submitted. REW provided code for the phenotype and genetic instrument generation of the lifetime smoking score, and reviewed and revised the manuscript and approved the final version of the manuscript as submitted.

## SUPPORTING INFORMATION CAPTIONS

**S1 Text. Supplementary text, tables and figures.**

**S1 File. Results from smoking heaviness MR-pheWAS.**

